# Development of a multivariate clinical prediction model for the diagnosis of mild stroke/TIA in physician first-contact patient settings

**DOI:** 10.1101/089227

**Authors:** Maximilian B. Bibok, Andrew M. Penn, Mary L. Lesperance, P. Stat, Kristine Votova, Robert Balshaw

## Abstract

**OBJECTIVE:** To develop a clinical prediction model for diagnosing mild stroke/transient ischemic attack (TIA) in first-contact patient settings. DESIGN: Retrospective study design utilizing logistic regression modeling of patient clinical symptoms collected from patient chart histories and referral data. SETTING: Regional fast-track TIA clinic on Vancouver Island, Canada, accepting referrals from emergency departments (ED) and general practice (GP). PARTICIPANTS: Model development: 4187 ED and GP referred patients from 2008–2011 who were assessed at the TIA clinic. Temporal hold-out validation: 1953 ED and GP referred patients from 2012–2013 assessed at the same clinic. OUTCOMES: Diagnosis of mild stroke/TIA by clinic neurologists. RESULTS: 123 candidate predictors were assessed using univariate feature selection for inclusion in the model, and culminated in the selection of 50 clinical features. Post-hoc investigation of the selected predictors revealed 12 clinically relevant interaction terms. Model performance on the temporal hold-out validation set achieved a sensitivity/specificity of 71.8% / 72.8% using the ROC01 cutpoint (≥ 0.662), and an AUC of 79.9% (95% CI, 77.9%–81.9%). In comparison, the ABCD2 score (≥ 4) achieved a sensitivity/specificity of 70.4% / 54.5% and an AUC of 67.5% (95% CI, 65.2%–69.9%). The logistic regression model demonstrated good calibration on the hold-out set (β_0_ = −0.257); β_linear_ = 1.047).

**CONCLUSIONS:** The developed diagnostic model performs better than the ABCD2 score at diagnosing mild stroke/TIA on the basis of clinical symptoms. The model has the potential to replace the use of the prognostic ABCD2 score in diagnostic medical contexts in which the ABCD2 score is currently used, such as patient triage.

## Introduction

Stroke is one of the leading causes of death and long-term disability in North America.^1,2^ Many strokes are preceded by transient ischemic attacks (TIA), or “mini-strokes”.^3^ The continuum from TIA to ischemic stroke has been referred to as acute cerebrovascular syndrome.^4^ It is estimated that patients diagnosed with a TIA have a 10% risk of recurrent stroke within 90 days (5% within 2 days) if untreated, with 50% of recurrent strokes occurring within the first 48 hours.^1^ Early treatment is associated with better patient outcomes, and is the target of many stroke best practice guidelines.^1,2^

In response to the growing awareness of the importance of TIA management for stroke prevention, Canadian stroke best practices recommend that all patients suspected of a TIA be assessed by stroke specialists within 48 hours of symptom onset.^1,2^ TIA clinics are frequently overburdened by referral volumes making the urgent triage of such high-priority patients difficult. Successful adoption of such recommendations, therefore, rest on the ability of TIA clinics to more accurately triage patients.

Diagnosis of ACVS can be challenging for TIA clinic staff as ACVS phenotypes can vary widely between patients depending upon the brain region affected. Compounding this difficulty many noncerebrovascular conditions, such as acute migraine, share many of the same features as ACVS. Such noncerebrovascular conditions, or mimic conditions,^5^ make it difficult for clinic staff to presumptively diagnosis referrals in order to prioritize true ACVS referrals for clinic intake. Approximately 30–60% of referrals to specialized TIA/stroke units are ultimately diagnosed as mimic conditions.^6–12^ Mimic referrals overburden the limited capacity and resources of specialized TIA/stroke units, and this increases patient wait times and delays the timely arrival of ACVS patients to these units. Assisting TIA clinic staff in differentiating ACVS from mimic conditions could serve as a first step toward improving TIA management.

Several studies have examined the viability of using the ABCD2 score to diagnosis ACVS.^8,9^ Originally developed as a prognostic tool for predicting risk of recurrent stroke after a TIA, the ABCD2 score has been found to be moderately diagnostic of ACVS.^8^ This is due in large part to the presence of several clinical features (e.g., age, blood pressure, clinical presentation) within the ABCD2 score that are also diagnostic of ACVS. Previous studies^9,13^ have reported low sensitivities (60.3–67%, cutpoint ≥ 4) for the score to diagnosis of ACVS and would appear to limit its clinical usefulness. This is unfortunate as approximately one-third of TIA clinics triage referrals on the basis of the ABCD2 score.^14^

Our research group has been working on a clinical prediction rule (CPR) to differentiate ACVS from mimic patients on the basis of patients’ history of presenting symptoms. Such a CPR would have value in the triaging of patients referred to TIA clinics. A deficiency of using the prognostic ABCD2 score as a diagnostic tool is that due to its design it contains no mimic predictors, as the diagnosis of ACVS has already been established prior to prognostic use. When used as a diagnostic tool the ABCD2 score structurally only provides a presumptive diagnosis of ACVS; all other diagnoses are implicitly defined as not ACVS. In contrast, we approached the problem of ACVS recognition as a matter of differential diagnosis. We appreciated that adopting such a framework will necessarily result in a model containing a large number of clinical features. This design decision runs counter to the principles of simplicity and brevity underlying most clinical support tools currently used in clinical practice. Instead, our aim when constructing our model was to focus exclusively on the task of differentiating ACVS and mimic conditions. To the best of our knowledge, no studies currently exist in the literature that approaches the issue of ACVS recognition as we have.

To develop our CPR we will assemble a dataset containing key clinical predictors previously identified in the literature and informed by current best practice as symptoms of ACVS and mimic conditions. From the dataset, we will use logistic regression modelling guided by clinical insights to construct our CPR. Finally, we will validate the model on a temporal hold-out dataset. The contributions of our study to the ACVS literature are to develop a CPR to specifically differentiating ACVS and mimic conditions

## Methods

### Settings and Participants

We utilized a retrospective study design to construct our study dataset. Patient clinical records from the Stroke Rapid Assessment Unit (SRAU), Victoria, B.C., from 2008–2013 were used to construct the dataset. The SRAU is a specialized outpatient stroke unit servicing most of the Vancouver Island (pop. 759,366). The SRAU receives referrals from emergency departments, general practice, and specialists (e.g., ophthalmologists). Median wait-time from referral to unit arrival was 4.397 days (IQR = 1.864–9.523). Only patient first time assessments at the SRAU during the time period were included in the dataset (N = 7823). The Health Research Ethics Board of Island Health (one of five provincial health authorities in British Columbia) approved this study. Funding for this study was provided by the Heart and Stroke Foundation, Genome British Columbia, Genome Canada, and Canadian Institutes of Health Research.

### Data Preparation and Missing Data

The dataset consisted of patients’ reported event histories, risk factors (age, sex, hypertension, diabetes, etc.), ABCD2 scores (derived by adding patient diabetes status to the ABCD score recorded in the patients’ charts), and final diagnoses. Event histories consisted of two free text fields (chief complaint and history of presenting illness). These two free text fields (henceforth, history fields) consist of descriptions of the patients’ events as recounted by the patients. Effort is made by SRAU stroke nurses to record the patients’ histories using the same language and words provided by the patient. Patient histories are recorded prior to a complete neurological assessment and final diagnosis by SRAU neurologists.

History fields were codified by a simple text-mining procedure that used pattern matching, in conjunction with NegEx^15^ negation detection, to map keywords and phrases onto clinical symptoms. Processing of the history fields is described in full in Sedghi et al., 2015.^16^ The clinical symptoms that were codified were selected in advance of statistical analyzes. Symptoms to be codified were chosen that were either (a) well referenced in the ACVS literature (e.g., vertigo, unsteadiness), or (b) medical concepts that appeared in the patients’ history fields (i.e., data-driven symptoms selection; e.g., Electric sensation, Emotionally labile, etc.). Absence of keywords and phrases in records were coded as the absence of the symptom in the patients’ records.

The complete initial dataset contained a total of 121 variables (diagnosis, risk factors, ABCD2 score, and codified history fields). The only continuous variables in the dataset were age (years), and systolic and diastolic blood pressure (mmHg). All other predictors were coded as: absent/no = 0, present/yes = 1; for patient sex, female = 0 and male = 1; for ABCD2 = 0, 1, 2, 3, 4, 5, 6, 7, or missing.

Patient diagnoses were by unit neurologists during standard of care neurological assessment. The diagnosis field extracted from the patient records consisted of eight possible classifications: (a) Stroke, (b) Stroke–Probable, (c) Stroke–Possible, (d) TIA, (e) TIA–Probable, (f) TIA–Possible, (g) Mimic, (h) Other, and (i) Unknown. Mimic conditions^5^ have similar clinical presentations/symptoms as TIA/mild Stroke, but are non-cerebrovascular in origin (e.g., migraine, seizure). The Other classification represents either non-ischemic strokes (e.g., hemorrhagic stroke) or non-cerebral ischemic events (e.g., cranial nerve ischemia). Unknown represents situations in which a diagnosis could not be determined (e.g., not yet diagnosed). Cases in the dataset were restricted to patients for whom either a clinical or radiological diagnosis (e.g., brain imaging) could be determined. Cases with classifications of Other or Unknown were omitted from analysis due to the ambiguity surrounding the diagnosis (see Figure 1). The remaining classifications were dichotomized as either ACVS (incl. Stroke, Stroke-Probable, Stroke-Possible, TIA, TIA-Probable, TIA-Possible) or Mimic for the purposes of these analyzes.

**Figure 1.**
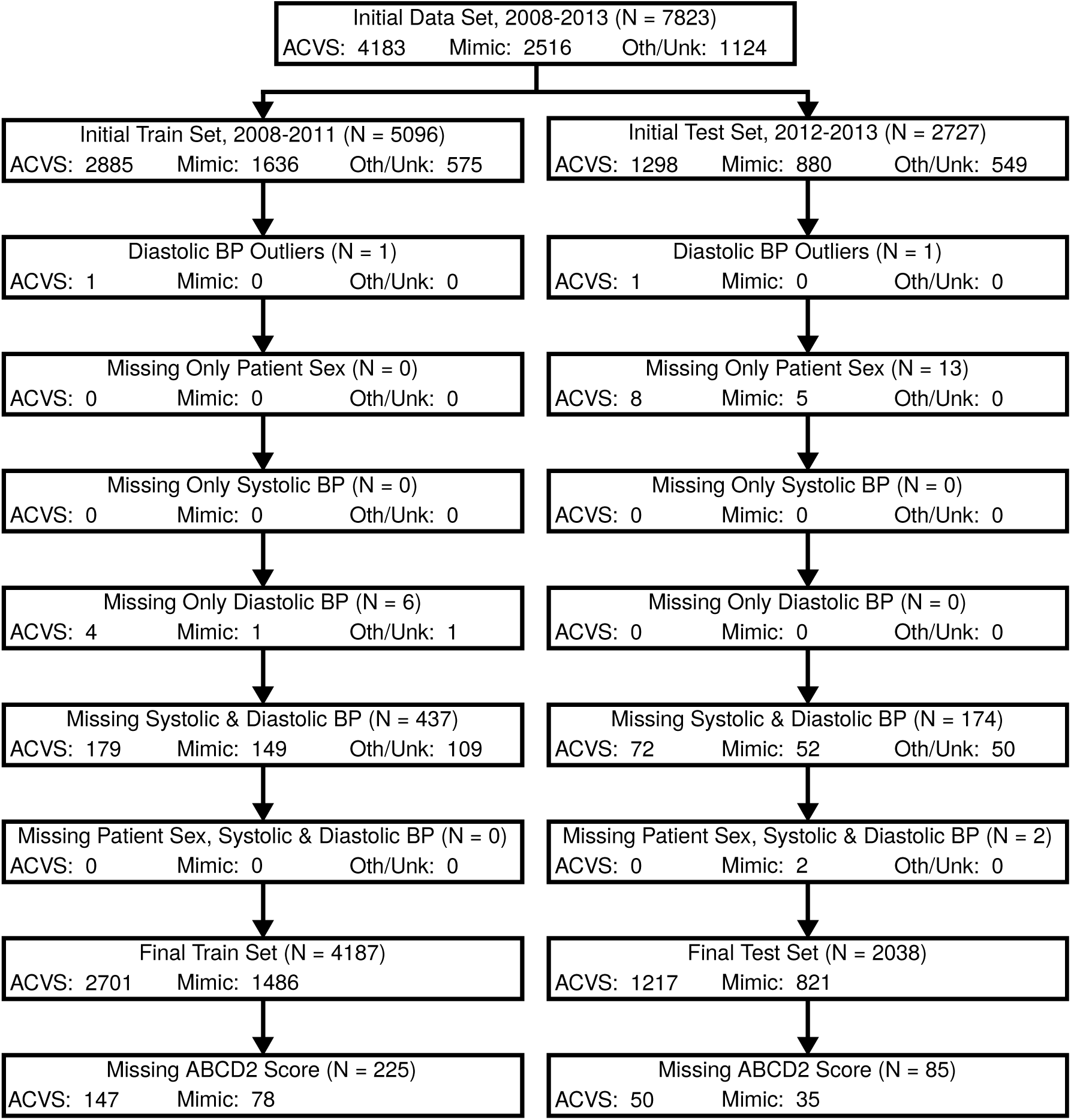
Participant flow diagram detailing missing data by variable and data set.

Prior to analysis, the variables comprising the initial dataset were reviewed for possible clinical redundancies. After review it was decided that several of the concepts be grouped together as composite variables and added to the dataset. Syncope and LOC (Loss of Consciousness) were combined with a logical OR operator to create Syncope or Loss of consciousness (LOC)* (henceforth, an asterisk (*) denotes a composite variable); i.e., Syncope or Loss of Consciousness (LOC)* is coded as present if the patient reported either Syncope or Loss of Consciousness, or both. Similarly, the variables Anxiety and Stress were combined to create the single variable Anxiety or stress*. The variables Visual field, Left visual field, Left eye visual field, Right visual field, and Right eye visual field were combined to create Visual field deficit (either side)*. Finally, Involuntary movement, Shaking, and Tremor were combined to create a single composite variable, Involuntary movement, shaking, or tremor*.

### Treatment of Missing Data

Prior to analysis, the data were examined for missing data and outliers (see Figure 1). Missing values were defined as any NULL values observed in the data extracted from the SRAU EMR database. The only outliers observed were two instances in which a diastolic blood pressure was recorded in excess of 700 mmHg. The only variables containing missing data were patient sex, systolic and diastolic blood pressure, and ABCD2 score. For these variables we assumed values were either not measured or recorded and thus were missing completely at random. With the exception of the ABCD2 score, listwise deletion was employed for the other variables prior to data analysis. In the case of the ABCD2 score, listwise deletion was applied to the data after feature selection was performed.

### Training and Test Datasets

The full clinical dataset (N = 7823) was divided into training (2008–2011, N = 5096) and test (2012–2013, N = 2727) datasets. Figure 1 provides a summary of how patients from the training and test datasets were excluded due to missing data and diagnosis.

Inter-rater reliability was conducted on the training dataset to compare the accuracy of the text-mining procedure with that of manual coding of patient history fields. Due to the size of the training dataset (N = 5096) only 1% (N = 52) of the total cases were analyzed, with random case selection equally distributed across the four years (2008–2011, N = 13 cases/year). For each of the clinical variables coded from the history fields Gwet’s AC1 was computed.^17^ Median Gwet’s AC1 was 0.98 (IQR = 0.957–1).

Table 1 displays the demographic characteristics of the final training (2008–2011, N = 4187) and test (2012–2013, N = 2038) datasets after missing data was addressed, with the exception of the ABCD2 score.

**Table 1.** Demographic characteristics of training and test data.

|  |  | Training Set | Test Set | T and chi-square |
| --- | --- | --- | --- | --- |
|  | Levels | (2008–2011) | (2012–2013) | homogeneity test |
|  |  |  |  | p values |
| N |  | 4187 | 2038 |  |
| Patient Age, mean (sd) |  | 68.97 (13.81) | 68.89 (13.83) | 0.825 |
| Male, N (%) |  | 2073 (49.5) | 1040 (51.0) | 0.272 |
| Diagnosis of ACVS, N (%) |  | 2701 (64.5) | 1217 (59.7) | <0.001 |
| CTA Completed, N (%) |  | 933 (22.3) | 862 (42.3) | <0.001 |
| MRI Completed, N (%) |  | 791 (18.9) | 393 (19.3) | 0.737 |
| ABCD2, N (%) | 0 | 48 (1.1) | 21 (1.0) | 0.151 |
|  | 1 | 174 (4.2) | 114 (5.6) |  |
|  | 2 | 495 (11.8) | 256 (12.6) |  |
|  | 3 | 813 (19.4) | 383 (18.8) |  |
|  | 4 | 1014 (24.2) | 509 (25.0) |  |
|  | 5 | 786 (18.8) | 367 (18.0) |  |
|  | 6 | 545 (13.0) | 260 (12.8) |  |
|  | 7 | 87 (2.1) | 43 (2.1) |  |
|  | Missing | 225 (5.4) | 85 (4.2) |  |
| Systolic BP, mean (sd) |  | 140.94 (21.75) | 143.56 (22.83) | <0.001 |
| Diastolic BP, mean (sd) |  | 76.95 (10.88) | 79.44 (11.11) | <0.001 |
| Hypertension, N (%) |  | 2503 (59.8) | 1231 (60.4) | 0.658 |
| Hyperlipidaemia, N (%) |  | 1754 (41.9) | 864 (42.4) | 0.726 |
| Atrial Fibrillation, N (%) |  | 503 (12.0) | 245 (12.0) | 1.000 |
| Diabetes, N (%) |  | 737 (17.6) | 382 (18.7) | 0.287 |
| Smoking, N (%) |  | 537 (12.8) | 226 (11.1) | 0.055 |

### Feature Selection

The process of feature selection was conducted on the final training dataset (N = 4187). Each feature was considered as a univariate predictor. Feature selection was filter based^18^ and utilized a statistical association in conjunction with clinical expert review.^18–20^ For each of the 123 candidate predictors (i.e., 121 variables in the dataset, less the diagnosis and ABCD2 score, plus the additional 4 composite variables, i.e., 121 − 2 + 4 = 123) the bivariate association with the diagnosis variable was computed. Odds ratios were computed for each predictor by regressing each variable independently on the diagnosis using a logistic regression model with the variable under consideration as the sole predictor. The Wald p-value was used as the statistical test of significance. Alpha was set at 0.05 for each candidate predictor. No effort was made to control for multiple comparisons;^21–23^ that is, the decision as to a predictor’s spuriousness was decided upon the basis of domain knowledge (i.e., support for the predictor in the ACVS literature).

Feature selection followed two stages. In the first stage, variables with a significant p-value value (α = 0.05) would be preferentially (i.e., favourably) reviewed for selection on the basis of their clinical relevance. Two criteria disqualified significant variables from selection: (a) the variable was clinically ill-defined (i.e., non-descriptive), and (b) the variable clinically overlapped another significant variable that was better defined and clinically broader in application.

In the second stage, variables with non-significant associations with the diagnosis were examined. Only two criteria were used to qualify these variables for selection. First, non-significant variables could be selected to complete a related set of clinically relevant variables if other members of the set were statistically significant; this review resulted in the inclusion of only the variables associated with numbness and weakness. These variable sets cover multiple body regions (left and right side face, arm, and leg, along with any overall indication of the condition, numbness or weakness, respectively). If one body region was significant, the other regions were selected. The second criterion was that non-significant variables could be included if their clinical relevance was already well established in the literature, and the exclusion of the variable would be clinically questionable.

### Model Construction

From the set of features chosen through this two-stage process, our next step was to include all selected features in a logistic regression model. Continuous predictors were standardized before model fitting using the following centers/standard deviations so as to allow for comparisons with the ABCD2 score: (a) age: 60 years of age / 10 years SD; (b) systolic blood pressure: 140 mmHg / 10 mmHg SD; and (c) diastolic blood pressure: 90 mmHg / 10 mmHg SD. Binary predictors were not standardized. Post-hoc interaction terms were then evaluated to explore how clinically meaningful interactions influenced model fit. Evaluation of interaction terms was on the basis of both statistical significance and clinical relevance. The full clinical model then comprised all selected variables plus a small number of clinically meaningful and significant interactions.

For the full clinical model three different cutpoints were calculated on the training dataset (2008–2011): (a) maximum efficiency^24^—the cutpoint that maximizes accuracy; (b) maximum Kappa^24,25^—the cutpoint that maximizes Kappa (i.e., inter-rater agreement (Cohen’s Kappa) between a model’s predictions and the true class labels); and (c) ROC01^25^—the cutpoint that minimizes the distance between point (0,1) on the ROC plot and the ROC curves.

After fitting of the full clinical model was completed we created an extreme phenotype variant of the model. In this variant, patients with clinical diagnoses of either TIA-Possible or Stroke-Possible were excluded from the training dataset, and the model was refitted (N = 3722). This model was created to examine the influence of these edge cases on the model’s performance. Cutpoints for this model were calculated as previously described.

Once models were examined for performance, final models for the full clinical (N = 6225) and extreme phenotype (N = 5510) models were constructed by refitting the models on the combined test and training sets, and cutpoints determined.

Although we did not conduct a formal power analysis, we considered the large number of records in the training set (N = 4187) as sufficient to pursue a multivariate model. A commonly used guideline^26^ indicates that an effective sample size of 10 ‘events’ per parameter is adequate for a logistic regression model. Our training set achieves 1486 ‘events’ (i.e., the smaller of the number of ACVS and the number of Mimics).

### Performance Evaluation

Performance of the full clinical model in differentiating ACVS patients from mimics was assessed on both the training dataset (in-sample performance) and the test dataset (out-of-sample performance). The test dataset was not used in the feature selection process so as to ensure an unbiased estimate of the full clinical model’s performance. Performance of the ABCD2 score for each dataset was also calculated so as to better contextualize the performance of the full clinical model. To evaluate the apparent performance of the full clinical model on the training set, cases with missing ABCD2 scores were removed after model fitting was completed. This was done to permit a direct comparison of the model’s performance with that of the ABCD2 score on the same cases. For the test dataset, cases with missing ABCD2 scores were removed before analysis. For the full clinical model, predicted probabilities were used to access model performance; for the ABCD2 score raw scores (range 0–7) were used.

Decision curve analysis (DCA) was conducted to assess the clinical utility of the full clinical model and nd the ABCD2 score.^27–29^ Although traditional measures of discrimination (e.g., sensitivity, C-statistic, etc.) and calibration assess model performance, they cannot indicate if a model is actually clinically useful.^30,31^ DCA attempts to quantify clinical utility using the concept of net benefit. Net benefit represents the number of true positives predicted by a model adjusted for by the number of false positives that have been weighted by the relative consequences (harm) of an unnecessary treatment relative to a missed treatment (i.e., *p* / (1 − *p*), where *p* is closely related to the concept of positive predictive value).^27,30,32^ Net benefit can also be understood as the gain in true positive cases detected by a model at a specific cutpoint, adjusted to be relative to a baseline “model” of assuming all cases to be negative (i.e., a “no treatment” or “none” model with no false positives).^27^

Analyzes were completed using the OptimalCutpoints (v1.1.3),^25^ PredictABEL (v1.2.2),^33^ ROCR (v1.0.7),^34^ pROC (v1.8),^35^ Hmisc (v3.17.4),^36^ rms (v4.5.0),^37^ immer (v0.5.0),^38^ dplyr (v0.5.0),^39^ and ggplot2 (v2.1.0)^40^ libraries in the R statistical language (v3.3.1).^41^

## Results

### Feature Selection

Feature selection results can be found in the Supplement. Table S1 displays the 123 candidate variables, along with their clinically related conditions, bivariate associations with the diagnosis (ACVS vs. Mimic), sorted by logistic regression Wald *p* value, and final selection decision. Odds ratios could not be calculated for two variables owing to low frequencies in the training set: Pallor (N = 3) and Collapse (N = 0).

During first stage selection (*p* < 0.05), 74 candidate variables were identified for review. Of these variables, 44 were retained, 19 were deemed clinically non-descriptive, 7 were constituents of the composite variables, and 4 were redundant with broader clinical concepts. Variables that were included as constituents of the composite variables (N = 7) were automatically excluded if the composite variable was significant. This occurred for the four composite variables, (a) Involuntary movement, shaking or tremor*, (b) Visual field deficit (either side)*, (c) Syncope or Loss of consciousness (LOC)*, and (d) Anxiety or stress*. The only clinically overlapping variables that were excluded were Face droop right, and Face droop left.

During second stage selection (*p* > 0.05), 47 candidate variables were identified for review. Of these variables, 35 were excluded, 4 were retained as clinically relevant, 6 were constituents of the composite variables, and 2 were retained to complete the numbness and weakness variable sets.

Of the 4 retained variables, three related to the visual domain, Curtain, Diplopia, and Vision loss, with the other two being Smoking and Eye droop (ptosis). The visual variables were retained as the visual domain has received relatively less attention in the ACVS literature compared to the motor and speech domain.^42^ Curtain represents a patient’s phenomenological (and indeed self-reported as such) experience of a “curtain” or “shade” descending over his or her field of view.^43^ This symptom is characteristic of transient monocular blindness (Amaurosis fugax). Vision loss represents the condition of homonymous hemianopsia, a symptom of posterior circulation strokes. Diplopia, or double vision, is another symptom of posterior circulation stroke. Smoking is known risk factor for stroke. Eye droop (ptosis) was retained due to its relation to Bell’s Palsy. Up to 75% of patients with Bell’s Palsy believe they have suffered a stroke,^44^ making Eye droop (ptosis) an important variable to include in the model.

To summarize, of the 123 candidate variables examined, a total of 50 variables were selected for inclusion in the final logistic regression model. Of these variables, 44 had a significant bivariate association (*p* < 0.05) with the diagnosis, and 6 were not significantly associated with the diagnosis but either were, (a) included to complete the numbness and weakness variable sets (N = 2), or (b) included on the basis of clinical relevance (N = 4).

In the logistic regression model fitting, twelve clinically relevant interaction terms based upon the selected predictors were included in the model. Table 2 displays the full clinical model and supporting references in the literature for the predictors in the model. Table 3 displays the extreme phenotype variant of the model.

**Table 2.** Full Clinical logistic regression model (Age, Systolic BP and Diastolic BP standardized). B=coefficient estimate, OR=odds ratio, CI=confidence interval.

| Variable | B | OR | 95% CI |
| --- | --- | --- | --- |
| Intercept | -0.489 | 0.613 | 0.476–0.789 |
| Patient age (years, center = 60, sd = 10)<sup>5,11,54–57</sup> | 0.301 | 1.351 | 1.248–1.463 |
| Patient sex<sup>5,7,11,57</sup> | 0.441 | 1.554 | 1.320–1.829 |
| History of hypertension<sup>5,11,57,58</sup> | 0.335 | 1.398 | 1.189–1.643 |
| History of hyperlipidaemia<sup>11,57</sup> | 0.121 | 1.128 | 0.964–1.321 |
| Smoking<sup>5,11,57</sup> | 0.470 | 1.600 | 1.266–2.023 |
| History of atrial fibrillation<sup>58</sup> | 0.224 | 1.251 | 0.977–1.602 |
| History of diabetes<sup>5,11,57</sup> | 0.082 | 1.086 | 0.885–1.332 |
| History of migraine<sup>54,57</sup> | -0.474 | 0.623 | 0.454–0.853 |
| Dizziness<sup>5,57,59–61</sup> | 0.006 | 1.006 | 0.817–1.239 |
| Headache<sup>5,54,55,62</sup> | -0.324 | 0.723 | 0.590–0.887 |
| Systolic BP (mmHg, center = 140, sd = 10)<sup>56</sup> | 0.055 | 1.056 | 1.006–1.109 |
| Diastolic BP (mmHg, center = 90, sd = 10)<sup>56</sup> | 0.026 | 1.026 | 0.934–1.127 |
| Face droop<sup>56,63,64</sup> | 0.673 | 1.960 | 1.431–2.685 |
| Weak face left<sup>56,63,64</sup> | -0.206 | 0.814 | 0.459–1.443 |
| Weak face right<sup>56,63,64</sup> | -0.700 | 0.497 | 0.277–0.892 |
| Weak arm left<sup>56,63,64</sup> | 0.470 | 1.600 | 1.137–2.251 |
| Weak arm right<sup>56,63,64</sup> | 0.568 | 1.765 | 1.233–2.528 |
| Weak leg left<sup>56,63</sup> | 0.277 | 1.319 | 0.864–2.014 |
| Weak leg right<sup>56,63</sup> | 0.420 | 1.523 | 0.948–2.446 |
| Weakness (motor, any indication)<sup>11,56,63</sup> | 0.426 | 1.532 | 1.247–1.881 |
| Numb face left<sup>62,63</sup> | 0.503 | 1.653 | 1.192–2.292 |
| Numb face right<sup>62,63</sup> | 0.220 | 1.246 | 0.847–1.832 |
| Numb arm left<sup>62,63</sup> | 0.205 | 1.227 | 0.915–1.646 |
| Numb arm right<sup>62,63</sup> | 0.165 | 1.179 | 0.837–1.661 |
| Numb leg left<sup>62,63</sup> | 0.498 | 1.646 | 1.100–2.462 |
| Numb leg right<sup>62,63</sup> | 0.442 | 1.556 | 0.950–2.548 |
| Numbness (any indication)<sup>11,62,63</sup> | -0.019 | 0.981 | 0.778–1.236 |
| Bilateral (numbness or weakness)<sup>5,60,65</sup> | -0.551 | 0.576 | 0.417–0.798 |
| Eye droop (ptosis)<sup>5,60</sup> | -0.030 | 0.971 | 0.432–2.180 |
| Visual field deficit (either side)*<sup>60,62,63</sup> | 0.366 | 1.442 | 1.018–2.041 |
| Neck pain<sup>55,60,66</sup> | -0.509 | 0.601 | 0.373–0.968 |
| Nausea<sup>5,60</sup> | -0.435 | 0.647 | 0.482–0.870 |
| Unsteadiness<sup>6,60,62,63</sup> | 0.256 | 1.292 | 1.074–1.553 |
| Curtain<sup>5,6,42,43</sup> | 0.411 | 1.509 | 0.986–2.308 |
| Syncope or Loss of consciousness (LOC)*<sup>5,11,12,42</sup> | -0.595 | 0.552 | 0.386–0.789 |
| Lightheaded<sup>5,11,12,61</sup> | 0.021 | 1.021 | 0.695–1.500 |
| Vertigo<sup>5,6,11,57,59,60,62</sup> | -0.871 | 0.419 | 0.294–0.596 |
| Confusion<sup>5,6,57,60,67</sup> | -0.585 | 0.557 | 0.452–0.687 |
| Amnesia<sup>5,11,42,60,67</sup> | -1.569 | 0.208 | 0.085–0.512 |
| Language disturbance<sup>56,62–64</sup> | 0.450 | 1.568 | 1.246–1.973 |
| Speech<sup>56,62–64</sup> | 0.769 | 2.157 | 1.767–2.633 |
| Positive visual disturbance<sup>5,6,42,54</sup> | -0.866 | 0.421 | 0.321–0.551 |
| Position (movement of head causes Sx)<sup>5,59–61</sup> | -0.407 | 0.665 | 0.270–1.638 |
| Diplopia<sup>5,6,54,60,62</sup> | 0.525 | 1.690 | 1.228–2.326 |
| Vision loss<sup>5,42,60,62,63</sup> | 0.686 | 1.986 | 1.370–2.879 |
| Concentration, Loss of<sup>5</sup> | -0.885 | 0.413 | 0.233–0.731 |
| Sudden onset (symptoms)<sup>5–7</sup> | 0.191 | 1.211 | 1.006–1.456 |
| Sensory march<sup>5,6,54</sup> | -0.623 | 0.536 | 0.317–0.908 |
| Involuntary movement, shaking, tremor*<sup>5,6,43,60,62,67</sup> | -0.712 | 0.491 | 0.338–0.713 |
| Anxiety or stress*<sup>5,11,12</sup> | -0.657 | 0.518 | 0.382–0.702 |
| Patient age (years, center = 60, sd = 10) × Concentration, Loss of | 0.354 | 1.424 | 0.968–2.095 |
| Patient age (years, center = 60, sd = 10) × Headache | 0.159 | 1.173 | 1.038–1.324 |
| Patient sex × Nausea | 0.239 | 1.270 | 0.815–1.980 |
| Patient sex × Position | -0.871 | 0.418 | 0.127–1.382 |
| Confusion × Amnesia<sup>67</sup> | 1.479 | 4.390 | 1.297–14.861 |
| Dizziness × Lightheaded | -0.764 | 0.466 | 0.285–0.761 |
| Headache × Lightheaded | 0.615 | 1.850 | 1.102–3.104 |
| Headache × Vertigo<sup>60</sup> | 0.674 | 1.962 | 1.109–3.469 |
| Neck Pain × Language Disturbance<sup>66</sup> | 1.325 | 3.763 | 1.015–13.950 |
| Lightheaded × Speech | -0.907 | 0.404 | 0.243–0.671 |
| Vertigo × Speech<sup>60</sup> | 0.786 | 2.195 | 1.126–4.277 |
| Face droop × Eye droop (ptosis)<sup>5,60</sup> | -1.129 | 0.323 | 0.100–1.047 |

**Table 3.** Extreme Phenotype logistic regression model (Age, Systolic BP and Diastolic BP standardized). B=coefficient estimate, OR=odds ratio, CI=confidence interval.

| Variable | B | OR | 95% CI |
| --- | --- | --- | --- |
| Intercept | -0.843 | 0.430 | 0.327–0.566 |
| Patient age (years, center = 60, sd = 10)<sup>5,11,54–57</sup> | 0.360 | 1.433 | 1.313–1.563 |
| Patient sex<sup>5,7,11,57</sup> | 0.555 | 1.743 | 1.463–2.075 |
| History of hypertension<sup>5,11,57,58</sup> | 0.343 | 1.410 | 1.183–1.680 |
| History of hyperlipidaemia<sup>11,57</sup> | 0.085 | 1.089 | 0.919–1.292 |
| Smoking<sup>5,11,57</sup> | 0.651 | 1.918 | 1.495–2.461 |
| History of atrial fibrillation<sup>58</sup> | 0.223 | 1.250 | 0.960–1.627 |
| History of diabetes<sup>5,11,57</sup> | 0.078 | 1.081 | 0.869–1.345 |
| History of migraine<sup>54,57</sup> | -0.597 | 0.550 | 0.383–0.791 |
| Dizziness<sup>5,57,59–61</sup> | -0.047 | 0.954 | 0.761–1.196 |
| Headache<sup>5,54,55,62</sup> | -0.371 | 0.690 | 0.551–0.865 |
| Systolic BP (mmHg, center = 140, sd = 10)<sup>56</sup> | 0.062 | 1.063 | 1.009–1.120 |
| Diastolic BP (mmHg, center = 90, sd = 10)<sup>56</sup> | 0.065 | 1.067 | 0.964–1.181 |
| Face droop<sup>56,63,64</sup> | 0.792 | 2.207 | 1.587–3.071 |
| Weak face left<sup>56,63,64</sup> | -0.041 | 0.960 | 0.520–1.772 |
| Weak face right<sup>56,63,64</sup> | -0.704 | 0.495 | 0.267–0.915 |
| Weak arm left<sup>56,63,64</sup> | 0.435 | 1.545 | 1.073–2.227 |
| Weak arm right<sup>56,63,64</sup> | 0.687 | 1.987 | 1.357–2.909 |
| Weak leg left<sup>56,63</sup> | 0.331 | 1.392 | 0.887–2.183 |
| Weak leg right<sup>56,63</sup> | 0.416 | 1.515 | 0.923–2.488 |
| Weakness (motor, any indication)<sup>11,56,63</sup> | 0.567 | 1.762 | 1.416–2.194 |
| Numb face left<sup>62,63</sup> | 0.485 | 1.625 | 1.139–2.317 |
| Numb face right<sup>62,63</sup> | 0.104 | 1.109 | 0.725–1.697 |
| Numb arm left<sup>62,63</sup> | 0.233 | 1.263 | 0.919–1.735 |
| Numb arm right<sup>62,63</sup> | 0.121 | 1.129 | 0.776–1.643 |
| Numb leg left<sup>62,63</sup> | 0.536 | 1.709 | 1.111–2.630 |
| Numb leg right<sup>62,63</sup> | 0.583 | 1.791 | 1.058–3.031 |
| Numbness (any indication)<sup>11,62,63</sup> | -0.071 | 0.932 | 0.726–1.196 |
| Bilateral (numbness or weakness)<sup>5,60,65</sup> | -0.660 | 0.517 | 0.360–0.742 |
| Eye droop (ptosis)<sup>5,60</sup> | 0.195 | 1.215 | 0.512–2.881 |
| Visual field deficit (either side)*<sup>60,62,63</sup> | 0.308 | 1.361 | 0.932–1.986 |
| Neck pain<sup>55,60,66</sup> | -0.481 | 0.618 | 0.368–1.039 |
| Nausea<sup>5,60</sup> | -0.621 | 0.538 | 0.381–0.759 |
| Unsteadiness<sup>6,60,62,63</sup> | 0.205 | 1.228 | 1.005–1.500 |
| Curtain<sup>5,6,42,43</sup> | 0.630 | 1.878 | 1.203–2.934 |
| Syncope or Loss of consciousness (LOC)*<sup>5,11,12,42</sup> | -0.714 | 0.490 | 0.330–0.727 |
| Lightheaded<sup>5,11,12,61</sup> | -0.050 | 0.951 | 0.626–1.446 |
| Vertigo<sup>5,6,11,57,59,60,62</sup> | -1.175 | 0.309 | 0.204–0.468 |
| Confusion<sup>5,6,57,60,67</sup> | -0.637 | 0.529 | 0.422–0.664 |
| Amnesia<sup>5,11,42,60,67</sup> | -2.010 | 0.134 | 0.039–0.463 |
| Language disturbance<sup>56,62–64</sup> | 0.495 | 1.641 | 1.286–2.093 |
| Speech<sup>56,62–64</sup> | 0.876 | 2.401 | 1.943–2.966 |
| Positive visual disturbance<sup>5,6,42,54</sup> | -0.834 | 0.434 | 0.323–0.584 |
| Position (movement of head causes Sx)<sup>5,59–61</sup> | 0.033 | 1.033 | 0.405–2.640 |
| Diplopia<sup>5,6,54,60,62</sup> | 0.605 | 1.831 | 1.305–2.570 |
| Vision loss<sup>5,42,60,62,63</sup> | 0.804 | 2.233 | 1.508–3.308 |
| Concentration, Loss of<sup>5</sup> | -0.942 | 0.390 | 0.200–0.761 |
| Sudden onset (symptoms)<sup>5–7</sup> | 0.248 | 1.282 | 1.051–1.563 |
| Sensory march<sup>5,6,54</sup> | -0.775 | 0.461 | 0.255–0.833 |
| Involuntary movement, shaking, tremor*<sup>5,6,43,60,62,67</sup> | -0.683 | 0.505 | 0.337–0.757 |
| Anxiety or stress*<sup>5,11,12</sup> | -0.936 | 0.392 | 0.275–0.559 |
| Patient age (years, center = 60, sd = 10) × Concentration, Loss of | 0.351 | 1.420 | 0.913–2.209 |
| Patient age (years, center = 60, sd = 10) × Headache | 0.200 | 1.222 | 1.065–1.402 |
| Patient sex × Nausea | 0.579 | 1.784 | 1.086–2.930 |
| Patient sex × Position | -1.219 | 0.296 | 0.083–1.053 |
| Confusion × Amnesia<sup>67</sup> | 1.930 | 6.890 | 1.466–32.378 |
| Dizziness × Lightheaded | -0.804 | 0.448 | 0.260–0.771 |
| Headache × Lightheaded | 0.693 | 1.999 | 1.122–3.560 |
| Headache × Vertigo<sup>60</sup> | 1.008 | 2.741 | 1.447–5.193 |
| Neck Pain × Language Disturbance<sup>66</sup> | 1.292 | 3.639 | 0.887–14.923 |
| Lightheaded × Speech | -1.057 | 0.348 | 0.199–0.608 |
| Vertigo × Speech<sup>60</sup> | 0.913 | 2.491 | 1.210–5.131 |
| Face droop × Eye droop (ptosis)<sup>5,60</sup> | -1.791 | 0.167 | 0.047–0.595 |

### Performance Evaluation

The full clinical model was fitted to the training data (N = 4187) and cutpoints for the model were determined. The predicted probability corresponding to each type of cutpoint were: (a) maximum efficiency: 0.516; (b) maximum Kappa: 0.59; and (c) ROC01: 0.662. For the ABCD2 score a cutpoint of ≥ 4 was used as this value has been suggested by a number of national stroke guidelines.^45,46^ These cutpoints were used to evaluate performance of the models on the test dataset.

The extreme phenotype variant for the full clinical model was fitted to the training data, less “Possible” diagnoses, (N = 3722) and cutpoints determined. The predicted probability corresponding to each type of cutpoint were: (a) maximum efficiency: 0.499; (b) maximum Kappa: 0.567; and (c) ROC01: 0.599.

Table 4 displays the performance measures of the full clinical model, extreme phenotype variant, and ABCD2 scores on both the training (re-substitution performance on training set) and test datasets, less patients with missing ABCD2 scores (N = 3962 and N = 1953, respectively).

**Table 4.** Performance measures (95% confidence interval) of models on training and test datasets.

|  | AUC | Sensitivity | Specificity | Accuracy | Cutpoint |
| --- | --- | --- | --- | --- | --- |
| Full Clinical Model (training) | 0.804 (0.790–0.818) | 0.873 (0.859–0.885) | 0.556 (0.530–0.582) | 0.760 (0.747–0.773) | MaxEfficiency: 0.516 |
| Full Clinical Model (training) | 0.804 (0.790–0.818) | 0.807 (0.792–0.822) | 0.645 (0.620–0.669) | 0.750 (0.736–0.763) | MaxKappa: 0.590 |
| Full Clinical Model (training) | 0.804 (0.790–0.818) | 0.717 (0.699–0.734) | 0.739 (0.716–0.762) | 0.725 (0.711–0.739) | ROC01: 0.662 |
| Full Clinical Model (test) | 0.799 (0.779–0.819) | 0.882 (0.862–0.899) | 0.545 (0.510–0.579) | 0.746 (0.726–0.765) | MaxEfficiency: 0.516 |
| Full Clinical Model (test) | 0.799 (0.779–0.819) | 0.806 (0.783–0.828) | 0.634 (0.599–0.667) | 0.737 (0.717–0.756) | MaxKappa: 0.590 |
| Full Clinical Model (test) | 0.799 (0.779–0.819) | 0.718 (0.692–0.743) | 0.728 (0.696–0.758) | 0.722 (0.702–0.741) | ROC01: 0.662 |
| Extreme Phenotype Variant (training) | 0.802 (0.788–0.816) | 0.813 (0.798–0.828) | 0.630 (0.604–0.655) | 0.748 (0.734–0.761) | MaxEfficiency: 0.499 |
| Extreme Phenotype Variant (training) | 0.802 (0.788–0.816) | 0.752 (0.735–0.769) | 0.707 (0.683–0.731) | 0.736 (0.722–0.750) | MaxKappa: 0.567 |
| Extreme Phenotype Variant (training) | 0.802 (0.788–0.816) | 0.720 (0.703–0.738) | 0.739 (0.716–0.762) | 0.727 (0.713–0.741) | ROC01: 0.599 |
| Extreme Phenotype Variant (test) | 0.799 (0.779–0.819) | 0.823 (0.801–0.844) | 0.622 (0.588–0.655) | 0.742 (0.723–0.761) | MaxEfficiency: 0.499 |
| Extreme Phenotype Variant (test) | 0.799 (0.779–0.819) | 0.757 (0.731–0.780) | 0.686 (0.652–0.717) | 0.728 (0.708–0.747) | MaxKappa: 0.567 |
| Extreme Phenotype Variant (test) | 0.799 (0.779–0.819) | 0.721 (0.694–0.746) | 0.719 (0.686–0.749) | 0.720 (0.700–0.739) | ROC01: 0.599 |
| ABCD2 (training) | 0.655 (0.638–0.672) | 0.691 (0.673–0.709) | 0.526 (0.500–0.552) | 0.633 (0.617–0.647) | Guideline: 4.000 |
| ABCD2 (test) | 0.675 (0.652–0.699) | 0.704 (0.677–0.729) | 0.545 (0.510–0.579) | 0.640 (0.618–0.661) | Guideline: 4.000 |

The scaled Brier score (0 = uninformative model, 1 = perfect prediction)^47^ for the full clinical model was 0.254. DeLong’s test of ROC curves^48^ indicated a significant difference between the full clinical model and ABCD2 score on both the training and test datasets, *p* < 0.001, respectively.

On the test dataset the scaled Brier score was 0.226. However, measures of the calibration line^49–51^ suggest that the full clinical model is well specified (β_linear_ = 1.047), but ‘calibration-in-the-large’^50^ indicates that predicted probabilities are systematically too high (β_0_ = −0.257). In other words, the model does a good job predicting observed probabilities across a broad range of predicted probabilities, though it consistently overestimates the probability of ACVS (Figure 2, solid curve below the line of equality). Figure 2 displays validation plots^47^ (graphical representation displaying both model discrimination and calibration) for the full clinical model on the training and test datasets. The distribution of predicted outcomes along the x-axis depicts the discriminate performance of the model to differentiate ACVS from mimic cases.^52^

**Figure 2.**
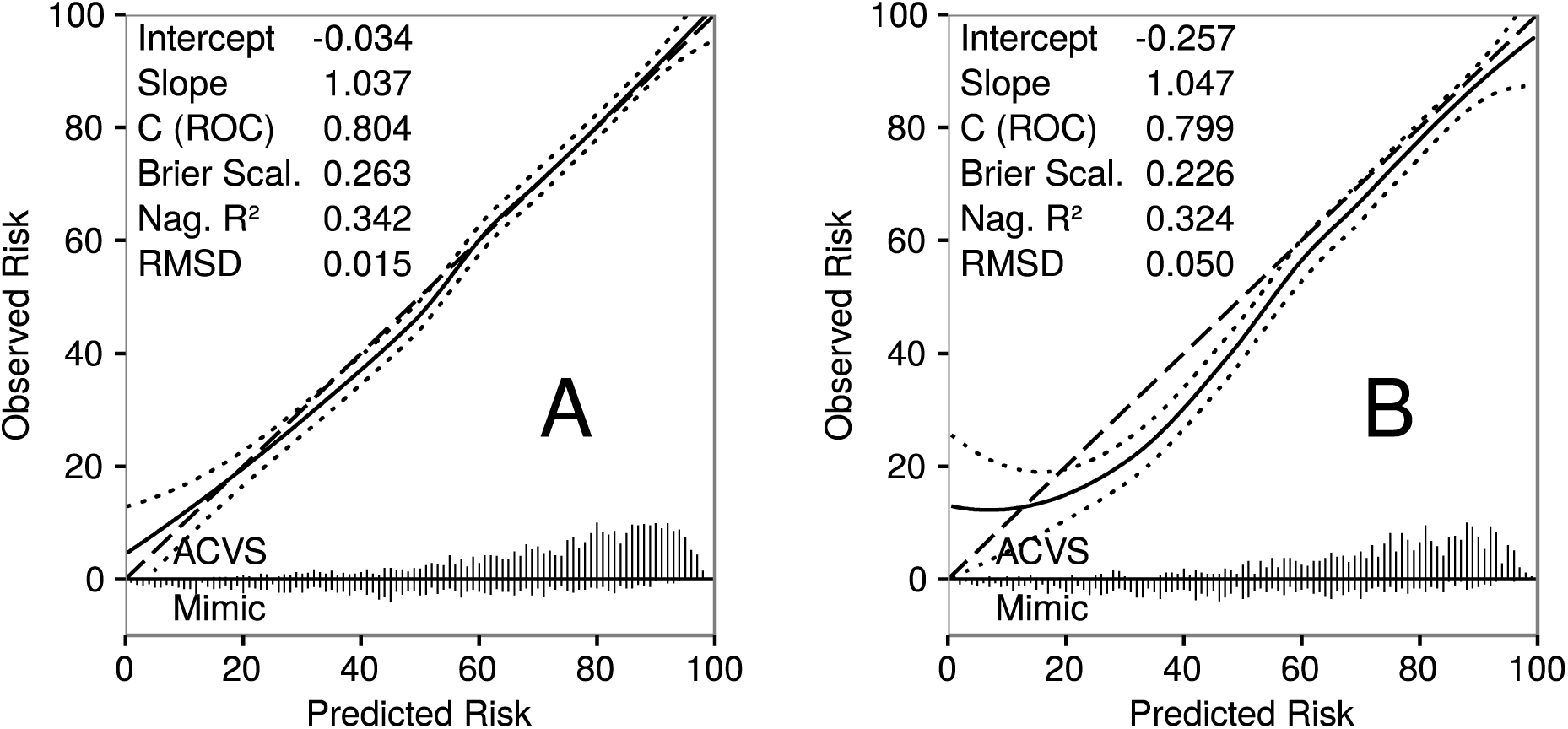
Validation plots of full clinical model on (A) training data, 2008–2011 (N = 3962), and (B) test data, 2012–2013 (N = 1953). RMSD=root mean standard deviation from line of equality.

Figure 3 displays decision curve analysis (DCA) plots for the full clinical model and ABCD2 score on the training and test datasets (ABCD2 scores were transformed to probabilities by way of logistic regression). On the test dataset the full clinical model achieved a net benefit of 0.214 at the empirical cutpoint determined on the training set, while the ABCD2 score achieved a net benefit of 0.118 at the empirical cutpoint of ≥ 4.

**Figure 3.**
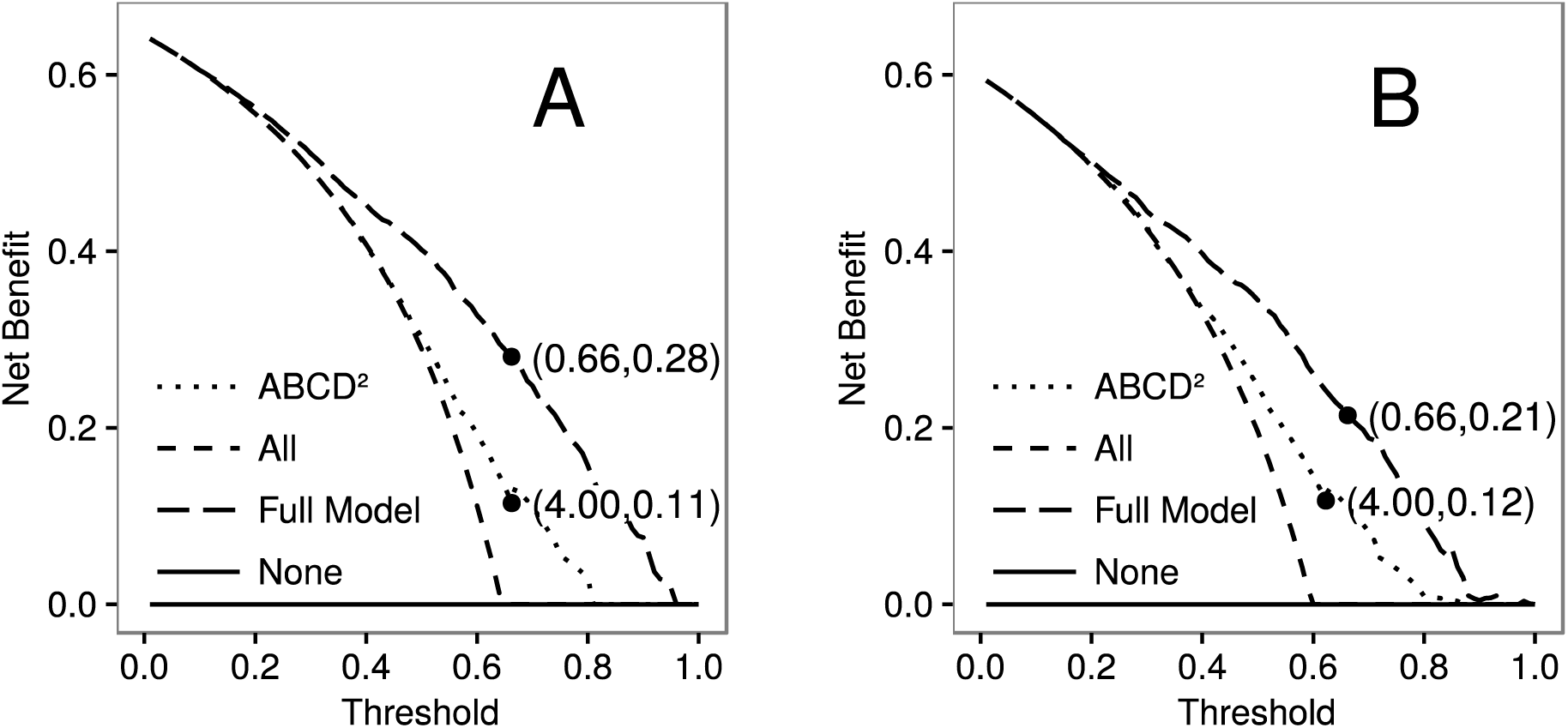
Decision curve analysis plots of full clinical model and ABCD2 scores on (A) training data, 2008–2011 (N = 3962), and (B) test data, 2012–2013 (N = 1953). Parentheses (cutpoint, net benefit).

### Final Model Fits

Tables S2 and S3 in the Supplement display the full clinical model and extreme phenotype variant model fit to the combined training and test datasets, respectively.

## Discussion

The goal of this study was to derive from patients’ reported event histories a clinically-informed model that can be used to optimally differentiate between ACVS and mimic conditions. Our full clinical model has demonstrated better predictive performance than the ABCD2 score. This increase in performance can be directly attributed to the extensive number of variables included in the full clinical model. The validation plots of the full clinical model on both the training and test sets suggest that the model is well specified and encompasses the major predictor variables of both ACVS and mimic conditions. Moreover, the AUC of the model on the test dataset (79.9%) is in keeping with theoretical upper limit for the AUC of a perfectly calibrated model.^53^ We interpret these results as evidence that the goals of the current study have been achieved.

The analysis of net benefit on the test dataset suggests that when performance is contextualized to a TIA clinic referral population (59.7% ACVS), our full clinical model achieves an increase in net benefit of 9.7% over the ABCD2 score. In the context of clinical practice (e.g., referral triage) the net benefit analysis suggests that our proposed clinical model might have greater clinical utility than the ABCD2 score as it currently used with a cutpoint of ≥ 4. Prior research has shown that a majority of specialized TIA clinics triage patient referrals by ABCD2 scores.^14^ As our model was developed on an equivalent population of patients it is likely that our model could be of value in improving upon existing TIA clinic triage practices.

A limitation of the current study is that the clinical practicality of the full clinical model is not immediately apparent. Given the goals of the study, the model is necessarily verbose with 50 main effects and 12 interaction terms. Moreover, logistic regression models are computationally complex and require the use of digital platforms to be usable as clinical prediction rules. In contrast, most clinical prediction rules tend to be terse and simple to complete, such as the ABCD2 score. Future work will need to determine if (a) such a large model can be made tractable for use by clinicians in real-world settings; or, conversely, (b) if a streamlined “bedside model” can be achieved by reducing the number of predictors and interactions in the model—in essence, simplifying the form of the model toward an additive point system like the ABCD2 score.

In conclusion, the results of the current study are encouraging in that they can be interpreted to suggest that a high degree of predictive ability to differentiate ACVS from mimic conditions can be attained on the basis of presenting clinical symptoms.

